# Precision timing with α-β oscillatory coupling: stopwatch or motor control?

**DOI:** 10.1101/591933

**Authors:** Tadeusz W. Kononowicz, Tillman Sander, Hedderik Van Rijn, Virginie van Wassenhove

## Abstract

Precise timing is crucial for many behaviors ranging from street crossing, conversational speech, to athletic performance. The precision of motor timing has been suggested to result from the strength of phase-amplitude coupling (PAC) between the phase of alpha oscillations (α, 8-12 Hz) and the power of beta activity (β, 14-30 Hz), herein referred to as α-β PAC. The amplitude of β oscillations has been proposed to code for temporally relevant information, and the locking of β power to the phase of α oscillations to maintain timing precision. Motor timing precision has at least two sources of variability: variability of timekeeping mechanism and variability of motor control. There is ambiguity to with of these two factors α-β PAC could be ascribed to. Whether α-β PAC indexes precision of internal timekeeping mechanisms like a stopwatch, or α-β PAC indexes motor control precision is unclear. To disentangle these two hypotheses, we tested how oscillatory coupling at different stages of time reproduction related to temporal precision. Human participants perceived, and subsequently reproduced, a time interval while magnetoencephalography was recorded. The data show a robust α-β PAC during both the encoding and the reproduction of a temporal interval, a pattern which could not be predicted for by the motor control account. Specifically, we found that timing precision resulted from the tradeoff between the strength of α-β PAC during the encoding and during the reproduction of intervals. We interpret these results as supporting evidence for the hypothesis that α-β PAC codes for precision of temporal representations in the human brain.

**Highlights:**

- Encoding and reproducing temporal intervals implicate α-β PAC.
- α-β PAC does not represent solely motor control.
- α-β PAC maintains the precision of temporal representations.

## Introduction

A precise understanding of the brain’s time-keeping precision is currently lacking. A number of human neuroimaging studies have started mapping our sense of elapsing time onto dynamical brain patterns recorded non-invasively with magnetoencephalography (MEG) and electroencephalography (EEG) (Macar, et al., 1999; Ng et al., 2011, Kononowicz & Van Rijn, 2011, 2015; Kulashekhar et al., 2015; Mento et al., 2013; Schlichting et al., 2018; Wiener et al., 2012, 2015; van Wassenhove & Lecoutre, 2015, for review, see Ng & Penney, 2014; Kononowicz et al., 2018). Most of these studies focused on neuronal indicators that scale linearly with the elapsed time (*e.g*., the contingent negative variation) thereby neglecting, the neuronal correlates of behavioral precision (i.e., variability). The precision of time-keeping mechanisms can bring novel insights to the neuronal mechanisms supporting timekeeping because it is free of other processes dynamically evolving in time that may be misleadingly interpreted to represent duration (*cf*. Gibbons & Rammsayer, 2004). Recent investigations have implicated beta (β) oscillations in time keeping process (Bartolo et al., 2015; Kononowicz & Van Rijn, 2015; Kononowicz et al, 2018; Kulashekhar et al., 2016; Wiener et al., 2018). Especially, during production of temporal intervals the power of β oscillations scales with the duration of produced interval (Kononowicz et al., 2018). However, consideration of only one frequency band could be oversimplifying for understanding of time keeping mechanisms. Based on the hypothesis that neural oscillations, and their coupling across time scales, are essential for the coding (Lisman & Jensen, 2013) and for the transmission (Fries, 2005) of information in the brain previous study asked whether the power of β oscillations may be regulated by the phase of alpha (α) oscillations during motor timing (Grabot, Kononowicz, et al., 2017). The strength of α-β phase-amplitude coupling (α-β PAC) was commensurate with the precision of the internally generated timed action such that narrower behavioral distributions were associated with stronger α-β PAC. These findings suggested that stronger α-β PAC allowed for a better maintenance, or precision, of temporal representations.

Note that β oscillations have been associated with both memory (Lundquist et al., 2016) and motor processes (Kilavik et al., 2013; Engel & Fries, 2010). The studies investigating the role of β power in time estimation do not provide a unified interpretation. The functional role of β power was originally described in motor tasks (Pfurscheller et al., 1996). β power is also associated with the control of motor commands (Swann et al., 2009). These studies support the notion that β power and motor control are highly associated (Brittain & Brown, 2014; Engel & Fries, 2010). Yet, β power effects were also reported in perceptual timing tasks in which no motor processes were required (Kulashekhar et al., 2016; Wiener et al., 2018), suggesting that β power could also influence non-motor processes. Our primary motivation was thus to make a distinction between motor and non-motor processes. On the basis of discrepancy between motor and non-motor processes in the interpretation of the role of β oscillations we derive two hypotheses: the motor control hypothesis, and working memory hypothesis. The first is involved with the precision of motor commands, the second one is involved with the precision of time keeping that necessarily has to involve working memory process (Gibbon et al., 1984). Working memory is a capacity for storing and manipulating information (Baddeley, 1992). In the context of interval timing it is uncertain how working memory operates (Gu et al., 2015; Matell et al. 2005).

Previous work linking β oscillations to timing has not been able to address which of these two processes was affected by coupling (Grabot, Kononowicz, et al., 2017), as time production involves a mixture of motor (motor implementation variability) and time-keeping processes (clock variability) (Keele et al., 1985).

In the study by Grabot, Kononowicz, et al., (2017), participants produced a short time interval by button presses: the first button press started the interval, and the second button press terminated the interval, with participants instructed to reproduce a duration as close as possible to the required target interval. In time production tasks (single interval), it’s difficult to distinguish between motor, perceptual, or central cognitive processes as all processes are involved in the production of that interval. To the contrary, in multi-duration time reproduction tasks, each trial starts with the presentation of a target duration, which then needs to be reproduced, typically via motor action. The separation in two task stages, namely, duration encoding and duration reproduction, thus provides a handle to investigate timing processes without the motor component that is not present in the encoding stage. This premise was supported by Baudouin et al., (2005) who showed that time production tends to be primarily associated with spontaneous tempo tasks, involving tapping at the preferred rate as regularly as possible, whereas time reproduction tend to be associated with working memory measurements, providing distinct cognitive blueprints in both tasks. Here, we focus on the difference between duration encoding and duration reproduction in the time reproduction task. Focusing on time reproduction as opposed to time production, ensures the differentiation of motor and working memory components. We ensure this separation to assess whether α-β PAC is linked with precision of motor control or with precision of working memory. The motor control hypothesis assumes that α-β PAC indexes precision with which motor commands and plans can be implemented, whereas the alternative hypothesis assumes that α-β PAC indexes the neural code for duration.

To differentiate motor control and working memory components we reanalyzed previously published data (Kononowicz et al., 2015) from a temporal reproduction task. As in every trial, participants were randomly presented with 2, 3, or 4 seconds intervals, which they had to reproduce, this dataset is well suited to investigate the contributions of α-β PAC during the encoding and the reproduction of durations. We set out to investigate if α-β PAC is present in both stages of time reproduction and whether α-β PAC in the encoding or reproduction is related to the precision of temporal durations.

## Methods

We reanalysed the data collected in Kononowicz et al., (2015) to investigate the role of α-β PAC during the encoding and reproduction of supra-second intervals.

### Participants

Eighteen students enrolled at the Humboldt, Freie or Technical University of Berlin with no self-reported hearing/vision loss or neurological pathology took part in the experiment in exchange for monetary compensation for participation. Written informed consent, as approved by the Ethical Committee Psychology of the University of Groningen, was provided by each participant before the experiment commenced. The data of two participants were discarded from the analyses: one participant fell asleep during the experiment, and the second one exhibited unreliable and spurious patterns of phase-amplitude coupling (PAC). The final sample comprised data of sixteen participants (all right handed, 8 males).

### Stimuli and Procedure

In a given trial, participants were presented with a to-be-memorized interval, called the Encoding Interval (EI, Fig. 1a) followed by the Reproduction Interval (RI), in which participants reproduced the EI.

**Figure 1.**
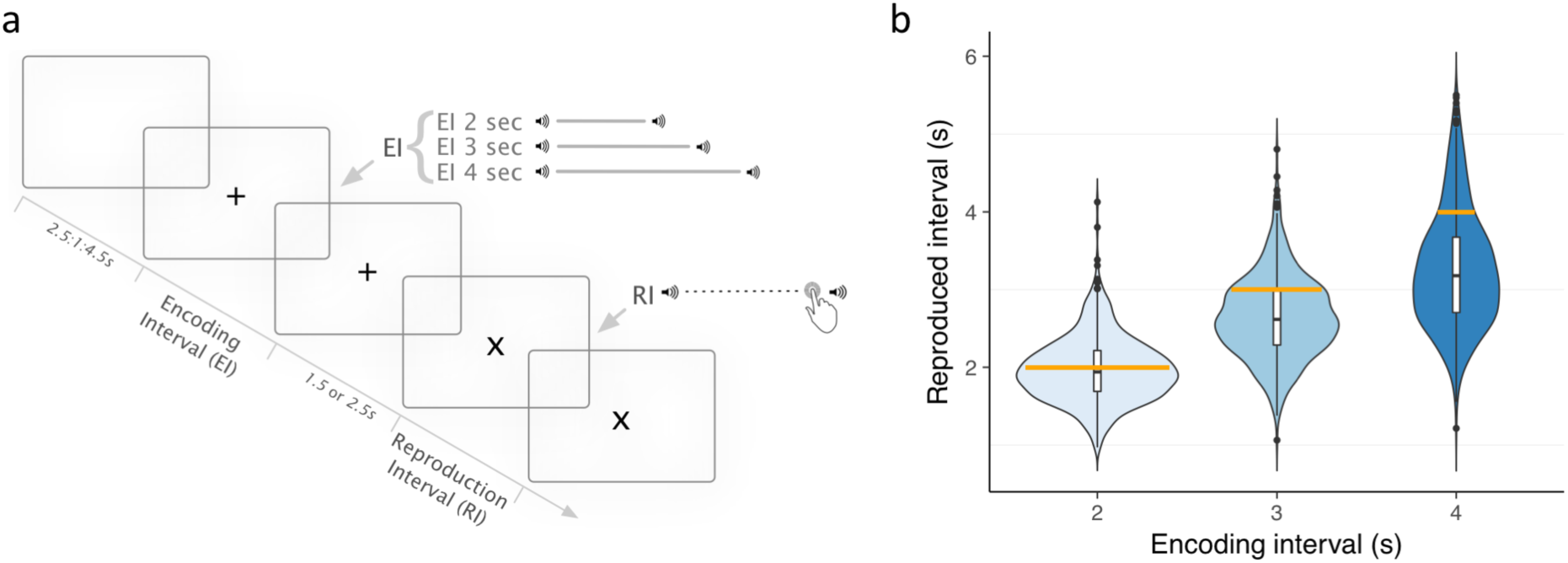
Task design and behavioral performance. a. Time course of the experimental trials in the time reproduction task. b. Probability density and box plots of time reproductions plotted for the three conditions separately. The yellow line demarcates the reference interval.

Each trial started with the presentation of a fixation cross “+”. After a randomly sampled inter-trial interval chosen among three possible values (2.5, 3.5 or 4.5 seconds (s)), the EI was presented as an empty interval delimited by two auditory tones (5 ms duration, 1 kHz, ~ 75 dB) separated by an interval of 2 s, 3 s or 4 s. The fixation cross “+” lasted for the entire time of the EI presentation, and up to 1.5 or 2.5 s after the EI, after which the reproduction interval (RI) started.

The start of the RI was indicated by the same tone as the EI, and with a change in fixation cross from “+” to “×”. Participants pressed the mouse button with their right hand when they thought the “×” was on the screen for the same amount of time as the “+” was. Pressing the response device button initiated the onset of a tone presentation, and the “×” disappeared from the screen. After an inter-trial interval randomly sampled out of 1.5 or 2.5 s, the next trial started. Each EI was presented 40 times yielding a total of 120 trials presented in 12 blocks of 10 trials each. Each block contained at least 3 repetitions of each EI pseudo-randomly presented. Between blocks, participants received adaptive feedback on their performance indicating how many trials were correct. The range of correct feedback was initially set to 20% deviation of the target interval and then dynamically adjusted by decreasing (−2.5%) or increasing (+0.5%) the range after each correct or incorrect trial, respectively. Participants were instructed to reproduce durations as accurately as possible and to maximize their number of correct trials in each block. Before the start of the MEG recordings, participants were presented with five practice trials. Stimuli were presented using a PC running Presentation software (Neurobehavioral Systems).

### MEG recordings

In a dimly-lit standard magnetically-shielded room (Ak3b, Vacuumschmelze, Hanau, Germany) located at the PTB Berlin, each participant laid in supine position with eyes open. Visual stimuli were presented on a screen via a projector located outside of the magnetically shielded room. Participants crossed their arms in front of their chest, as earlier work has shown this position to be most comfortable during longer recording sessions, and responded by clicking a button on a computer mouse, which was held in the right hand, located on their left upper body. Measurements were carried out with a Yokogawa MEG system (Yokogawa Electric Corporation, Japan) using 125 axial gradiometers with three reference magnetometers used for offline data denoising and ambient noise removal.

The head position with respect to the sensor helmet was measured using coils attached to the scalp at anatomical landmarks (nasion and preauricular points). The locations of the coils were digitized with respect to three anatomical landmarks with a 3D digitizer (Zebris, Isny, Germany).

The brief auditory bursts indicating the start and end of the intervals were presented binaurally via MEG-compatible tube earphones (Etymotic research, Elk Grove Village, USA) at sound levels set for each participant individually, at approximately 75 dB.

### Calculation and statistical assessment of phase-amplitude coupling based on Modulation Index

To prevent the influence of evoked responses in PAC estimation, we focused on the time segment starting from 0.4 s after the tone initiating the EI or the RI until the end of the EI duration (i.e., 2 s, 3 s, or 4 s). In the RI analysis, we focused on the time segment from 0.4 s following the stimulus onset up to 1.2 s in the 2 s condition. We chose 1.2 s, which was the lower tail of the RI distribution, to insure that the data were free of components associated with movement execution. For the same reason, in 3 s and 4 s reproduction intervals we focused on the period extending up to 2 s.

To calculate PAC, we split the data of each duration condition into temporal bins of similar length in order to equalize the number of time samples (given variability in produced duration lengths), and prevent erroneous assessment of PAC calculation (Dupré la Tour et al., 2017). To equate the amount of data, we used bins of 0.8 s long, starting at 0.4 s after the onset of the stimulus demarcating EI or RI onset. The number of bins depended on the duration condition and ranged from 1 to 4 bins for the 2 s to 4 s trials, respectively. Across all conditions, Bin1 ranged from 0.4 s to 1.2 s, Bin2 from 1.2 s to 2 s, Bin3 from 2.2 s to 3 s, and Bin4 from 3.2 s to 4 s.

The strength of PAC was assessed using the Modulation Index (MI; Tort et al., 2009; Dupré la Tour et al., 2017): raw data were band-pass filtered with a slow-frequency bandwidth of 2 Hz, and a high-frequency bandwidth of 20 Hz. The instantaneous amplitude of the high-frequency and the phase of the slow-frequency were extracted from the Hilbert transform applied to the epoched data. To assess whether the distributions diverged from uniformity, the Kullback-Leibler distance was computed, then normalized to provide an estimate of the strength of MI. The slow-frequency component ranged from 3 Hz to 15 Hz in steps of 0.5 Hz, and the high-frequency ranged from 14 Hz to 100 Hz in steps of 2 Hz. A comodulogram was computed for each MEG sensor, providing a full estimate of the MI over the full spectrum.

To assess the statistical significance of PAC at the individual level, the MI was compared to a surrogate distribution (n = 100) computed by randomly shifting the low frequency signal by a minimum of 0.3 s as was previously proposed by Tort et al. (2010). For each participant, 5 sensors with the highest Z-scores were selected and averaged. The resulting averages were plotted separately for the EI and the RI (Fig. 2). The resulting averaging of z-scores indicated the level of significance. Considering that Z-score can be converted to significance level, a Z-score of 4 corresponds to p = 0.00001.

**Figure 2.**
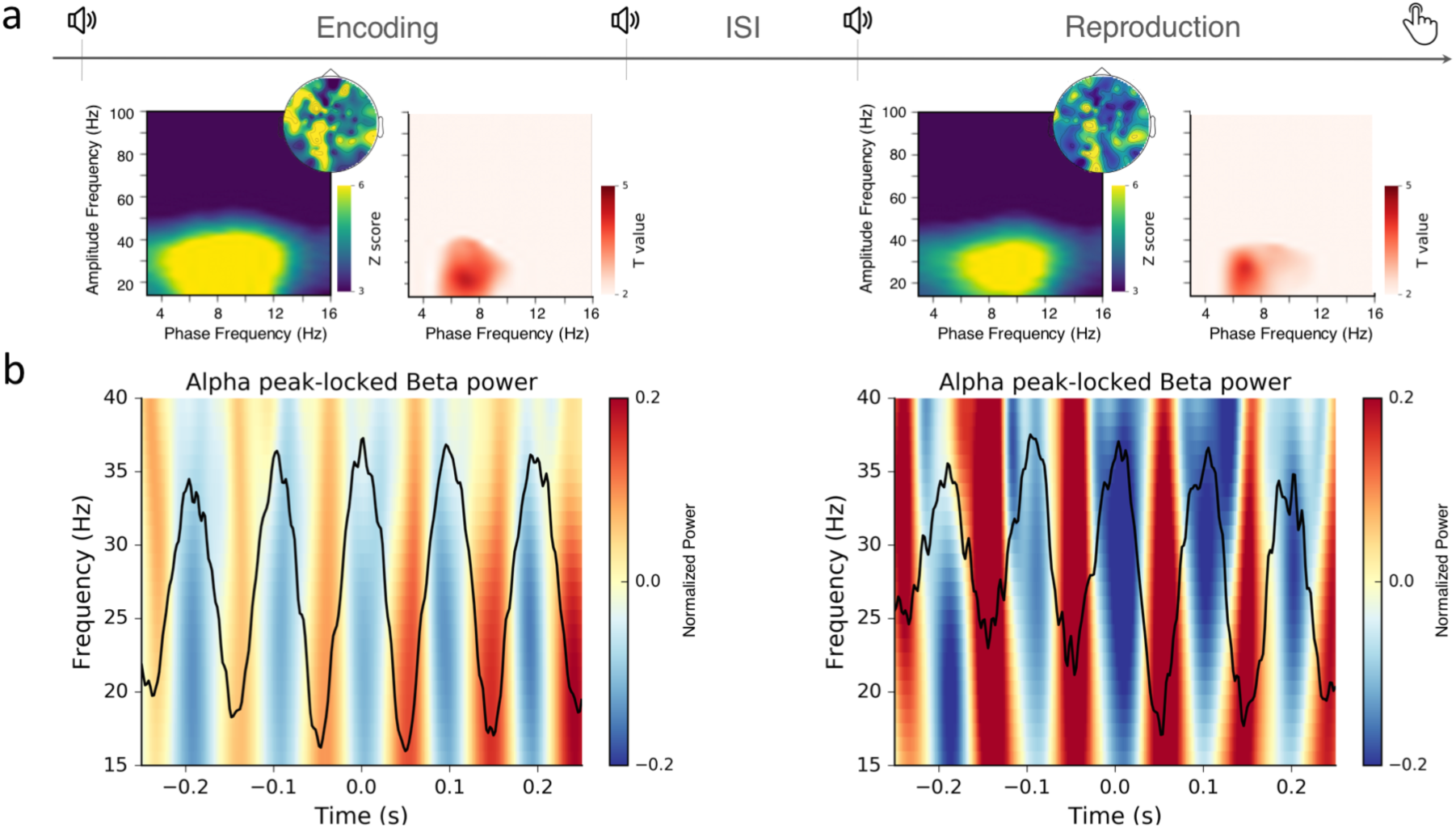
Robust phase-amplitude coupling in the encoding and in the reproduction of time intervals. a. Z-scored comodulograms (left) and t-scores resulting from permutation tests against the baseline interval in the encoding (left) and the reproduction (right) phases of the time reproduction task. b. Alpha peak-locking plot. The individual time series were locked on the peak of the α oscillations (black line). β Power (15-40Hz) was computed in the realigned data, normalized by subtracting its mean, and displayed as heatmaps. For illustration, we picked the MEG sensor showing maximal α-β MI for one representative participant.

To assess whether PAC was associated with the EI and with the RI, we compared the MI computed during the pre-trial baseline (from −0.8 s to 0 s prior to the first stimulus demarcating the EI onset) to the MI computed during the EI and the RI (Fig. 2). Statistical assessments of a difference using MI were performed using MNE’s python utility permutation_cluster_1samp_test with 1000 permutations, which implements the nonparametric statistical testing (Maris & Oostenveld, 2007). MI values arranged in a matrix were used as the input to the permutation test and subjects were used as observation samples (Fig. 2, red scale plots).

To assess stability of PAC, within the EI and RI, we compared consecutive bins within EI and RI. Similarly, to the analysis against the baseline, we used a cluster permutation test as in PAC on the whole range of computed MI values. Additionally, we also employed a second approach which focused on the α-β range of the MI matrix and used a t-test for repeated measures (ttest_rel as implemented in SciPy Python library).

### α peak-locking procedure for graphical representation of α-β PAC

To visualize α-β PAC in the time domain, we aligned α peak with the high frequency activity (here, β). Data were filtered with respect to the frequencies observed in the PAC analysis: as we observed a robust α-β PAC, we focused on illustrating how β power aligned with the phase of the α frequency. For this, we used a FIR filter implemented in the pactools Python package (v0.1, Dupré la Tour et al., 2017). α peaks were detected by localizing the maxima in the phase of the low frequency signal. The power of β was assessed by using a Hilbert transform to compute the instantaneous amplitude (the Hilbert analytical signal was multiplied with its conjugate and the real part was extracted). The resulting plots are displayed in Fig 2b.

### Localization of PAC peak using jackknife procedure

The location of the MI peak was quantified using a jackknife procedure. First, 5 sensors exhibiting maximal α-β PAC were identified and averaged by individual. Each ‘jackknife sample’ was obtained by removing a single sample and averaging the remaining samples (n-1, Miller et al., 1998; Ulrich & Miller, 2001). The coordinates of amplitude frequency and phase frequency of the MI were obtained on the basis of jackknife samples. F values, obtained by running the ANOVA (AnovaRM as implemented in statsmodels Python library, v0.9) on jackknife samples, were corrected by dividing obtained F by (n-1)^2^.

### Regression analyses between precision and PAC

The sensor level analyses were performed using linear regression and model comparison to check whether factors other than PAC were needed to account for timing behavior. All statistical analyses were performed using R v3.3.2 statistical programming language (R Development Core 2008).

Similar to the previous within participant approach, MEG data were randomly split into 4 trial-based bins. The 4-bins split was chosen to guarantee a maximal number of within-subject samples for the regression analyses while preserving a minimal number of samples for computation of the coefficients of variation (~10 samples per bin; van Belle, 2008). As for the previous analyses we selected 5 sensors with the highest MI for α–β PAC. To maximize PAC contribution to selected sensors, the sensors were selected per participant, condition (i.e., 2 s, 3 s, 4 s) and task stage (EI or RI). This ensured that each sample entered in the regression was effectively a product of most prominent PAC contributions.

The inverse coefficient of variation (invCV: mean duration production divided by the standard deviation in a set of approximately 10 trials) and the PAC comodulograms were computed for each data bin. Each outcome served as a single observation in regression model.

As we split the data per block and per individual, we used linear mixed-effects models (e.g., Pinheiro and Bates, 2000; Gelman and Hill, 2007) to account for multiple per subject observations in the data and samples dependencies. Linear mixed-effects models are regression models that model the data by taking into consideration multiple levels. Subjects and trial-based bins were random effects in the model, and were allowed to vary in their intercept. p values were calculated based on a Type-3 ANOVA with Satterthwaite approximation of degrees of freedom, using lmerTest package in R (Kuznetsowa et al., 2017). The mixed-effects models approach was combined with model comparisons that allow for selecting the best fitting model in a systematic manner.

Whenever Shapiro-Wilk normality test indicated a deviation from normality we transformed the data using using the Lambert W function. The Lambert W function provides an explicit inverse transformation, which removes heavy tails from the observed data (Georg, 2011, 2015). First, the data are transformed into a heavy-tailed form using log-likelihood decomposition. Subsequently the heavy tailed form is transformed back into a Gaussian distribution. All transformations were performed using LambertW R package.

## Results

Participants conformed to the task requirements as previously reported (Kononowicz et al., 2015; Fig. 1b). The distribution of time reproductions in the 2 s condition was centered at 2 s (mean = 1.98 s, SD = 0.42 s); the reproduction of 3 s (mean = 2.64 s, SD = 0.53 s) and 4 s (mean = 3.26 s, SD = 0.75 s) were shorter than the target duration (Fig. 1b). This pattern was consistent with Vierordt’s law and regression to the mean typically observed in time reproduction and estimation experiments (e.g., Jazayeri & Shadlen, 2010; Shi, Church & Meck, 2013; Petzschner et al., 2015; Polti et al, 2018).

### Robust α-β PAC during encoding and reproduction of temporal intervals

In previous work, we showed robust α-β PAC in temporal production focusing on MEG signals. Combined MEG and EEG data did not qualitatively changed the results (Grabot, Kononowicz et al., 2017). For simplicity, we will solely focus on MEG signals in the current paper.

First, we computed PAC over a broad spectrum of frequencies, and we collapsed the data across reproduced durations, participants, and sensors. Modulation Index (MI; Tort et al., 2009) quantifies phase-amplitude coupling by estimating a deviation of the amplitude distribution with respect to a certain phase from the uniform distribution, in a so-called phase-amplitude plot. We quantified the MI separately for the EI and the RI over the full spectrum to produce comodulograms. This allowed for identifying the peaks in the shuffled distribution of MI, and revealed significant peaks of α-β PAC (6-12 Hz for the carrier, and 15-40 Hz for the modulated frequency) in both EI and RI (see Fig. 2a for Z-scores and t-values). Although, we observed low frequency PAC ranging from 8 Hz to 12 Hz, in a previous report (Grabot, Kononowicz et al., 2017), here, the significant lower PAC frequencies appeared to extend to lower frequencies (Fig. 2a), with a center of gravity approximating 7 or 8 Hz. Therefore, we used an extended lower PAC frequency range (6-12 Hz) in all analyses.

We inspected the α-β PAC coupling regime by aligning β power to α oscillations in the time domain. The peak-locking procedure visualized the presence of α-β PAC and an example of realigned time series for a single participant is provided in Fig. 2b: for this participant, β power was highest at the trough of the α oscillation and lowest at the peak of the α oscillation. The comparison of each α-β PAC within each bin against the baseline interval converged with the permutation analysis of MI and visual inspection of peak-locked data. Each bin during encoding and reproduction significantly increased in the strength of α-β PAC as compared to baseline (Fig. 2a, the right panels for EI and RI).

The observation that α-β PAC during EI significantly increased as compared to baseline corroborated the prior working hypothesis that α-β PAC may plays a role in timing (Grabot, Kononowicz et al., 2017): considering that α-β PAC was present when participants were not required to perform motor actions, α-β PAC may go beyond precision in motor execution and motor preparation. A significant increase of α-β PAC was also observed during reproduction.

As stronger couplings in EI or RI could indicate predominant support of α-β PAC in one of the two stages of the task, we then contrasted MI in the EI and RI using a cluster permutation test. We found no evidence of significant α-β PAC difference between the EI and the RI (p > 0.1). Permutation tests were accompanied by Bayesian t test focused on α-β PAC (temporal bin1 in EI was tested against temporal bin1 in RI etc.). None of the tested comparisons showed BF exceeding 1.

### Stable α-β PAC during encoding and reproduction of temporal intervals

To delineate the temporal characteristics of α-β PAC, we assessed whether the magnitude of α-β PAC was stable or dynamic during the EI and RI (Fig. 3). For example, certain indices of interval timing, such as climbing neuronal activity measured as a gradual change of amplitude of Contingent Negative Variation (Walter, 1964) from interval onset until its offset, have been associated with dynamic processes related to duration perception (Boehm et al., 2014; Kononowicz & Van Rijn, 2014; Macar et al., 1999; Wiener et al., 2012). Along the same lines, we sought to investigate whether α-β PAC undergo similar dynamical changes within the EI or the RI, or whether it was stable within the timed interval.

**Figure 3.**
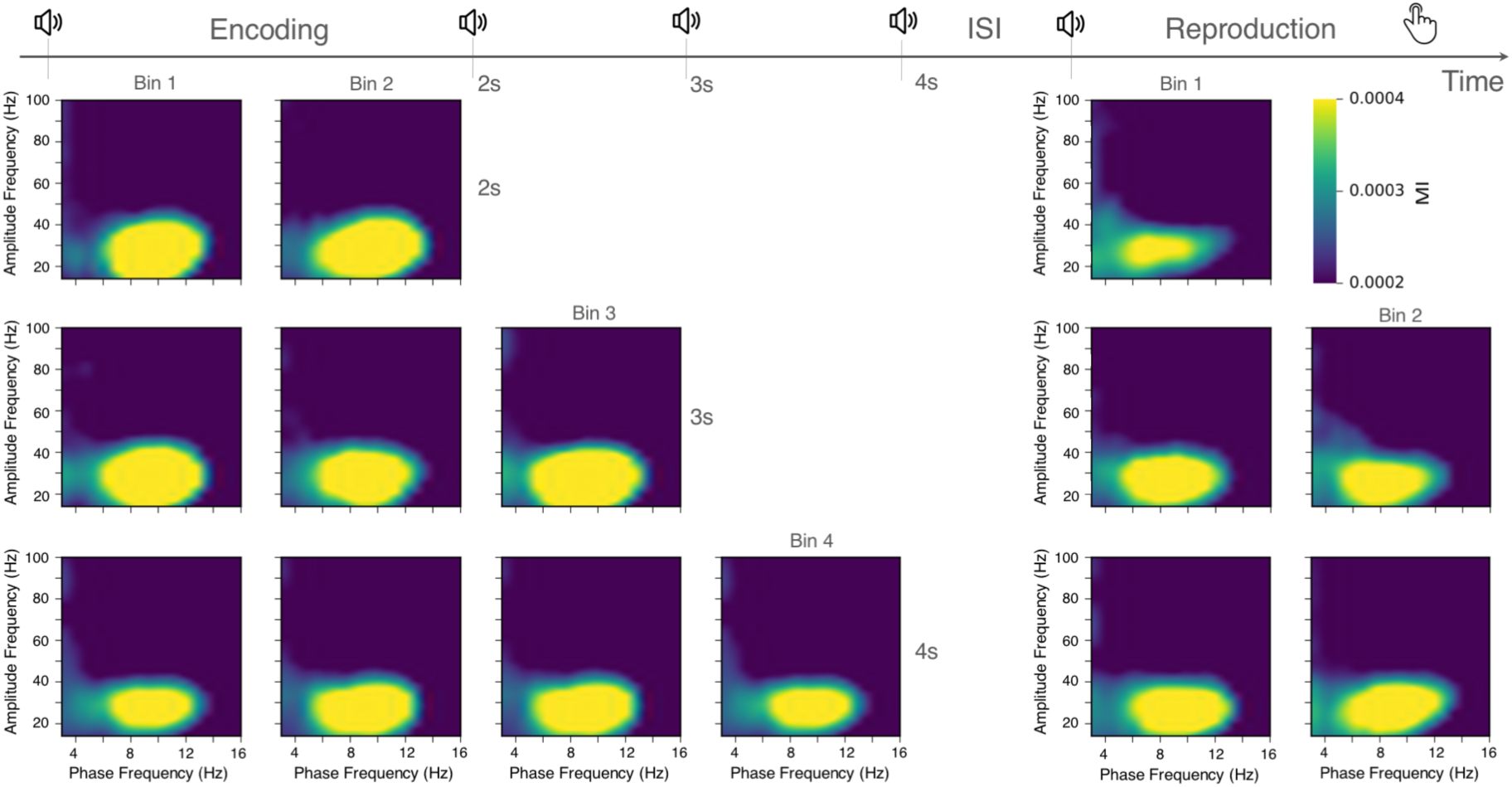
Stable strength of α-β PAC within trials. The comodulograms were inspected as a function of task stage and time bins within each experimental condition (columns) and reproduction length (rows). For each reported comodulogram, 5 sensors with maximal α-β PAC were identified per participant and grand-averaged across the sixteen participants.

According to the precision hypothesis, α-β PAC should be stable within the RI considering that it maintains the representation of a given duration. Alternatively, if α-β PAC supports dynamical features of timing related processes such as gradual integration of temporal information, we should systematically observe a covariation of α-β PAC with time, within a timed interval.

To test this, we used two different approaches. In a first approach, we focused on α-β PAC using collapsed values of 5 sensors with maximal MI per condition. We then performed a series of t-test comparisons between consecutive temporal bins, within EI, RI, and all duration conditions, also using Bayesian t test. None of the comparisons showed significant results (all p > 0.1; all BFs < 1). This series of comparison was not adjusted for multiple comparisons. In a second approach, we collapsed the data across all sensors and performed the permutation tests on a broad MI frequency spectrum, comparing consecutive bins in EI and RI. Again, none of the comparisons showed significant results (all p > 0.1). Hence, neither approach supported the notion of dynamical changes of α-β PAC during encoding, reproduction or across durations. This suggested a relatively stable α-β PAC within the timed interval, in line with the precision hypothesis.

### Stable α-β PAC across durations, encoding and reproduction

Considering that three durations were tested, we then questioned whether α-β PAC varied as a function of duration, during encoding and during reproduction. According to the precision hypothesis, one would not expect significant shifts of the PAC to be involved in the timing of different durations.

To explore the characteristics α-β PAC as a function of durations, we used the data depicted in Fig. 4: for each time bin, participant, EI, RI, and duration, we quantified the maximum coordinate of the MI using a jackknife method (Fig. 4). The obtained maximal amplitude of the high-frequency modulated by the maximal phase frequencies were entered into a two (stage: EI, RI) by three (durations: 2 s, 3 s, 4 s) repeated measures ANOVA. No significant (main or interaction) effects for amplitude frequency and phase frequency were found (all p > 0.1). Although Fig. 4 suggests a possible small effect of low frequency oscillation of PAC between the EI and the RI, a statistical test did not confirm this observation (F(1,16) = 1.7, p = 0.212). Hence, we found no evidence for a shift of PAC across EI, RI, or duration.

**Figure 4.**
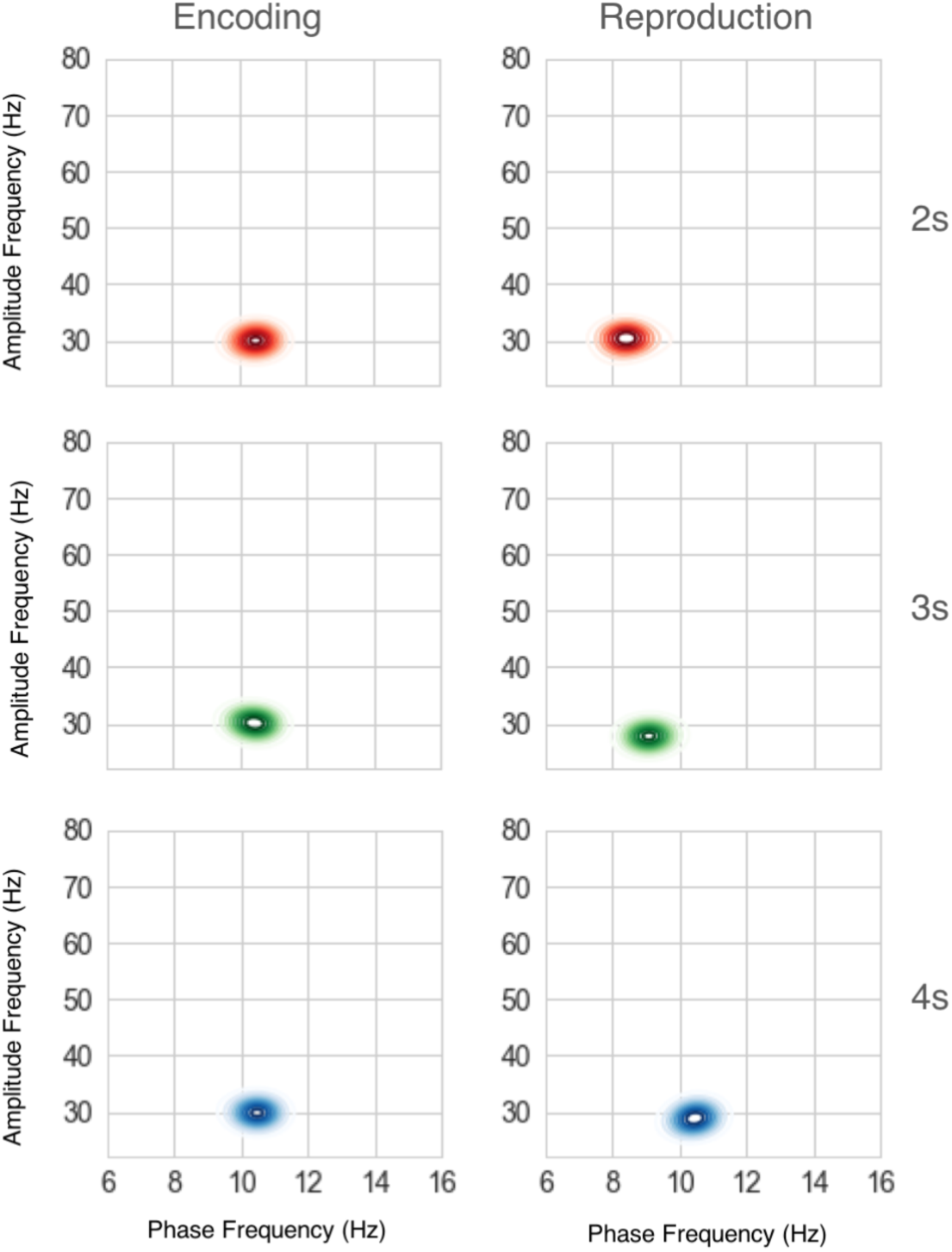
Stable peak of α-β PAC during trials and across reproduced durations. The location of the maximal amplitude frequency did not vary as a function of the reproduced time interval or task stage. The trend for the phase frequency to decrease during the reproduction interval did not reach significance. The individual data points were smoothed using 2D kernel density estimate as implemented in Seaborn python package.

### α-β PAC and timing precision

Different durations yielded different trial lengths. Given the absence of dynamic changes in α-β PAC within the trials, we focused regression analyses on the first and on the last bins for assessing the precision in EI, and on the first bin only to assess the precision in RI. We chose the first and the last bins under the assumption that the beginning and the end of EI, and the end of RI should represent the same cognitive steps irrespective of tested duration condition. Each observation was calculated using the average of 5 sensors with maximal α-β MI.

According to the precision hypothesis, α-β PAC may reflect the precision with which an endogenous timing goal may be maintained (Grabot, Kononowicz, et al., 2017). In our current reproduction paradigm, the encoding of the target duration was separated from its reproduction. Hence, the association between α-β PAC and timing precision could be expressed in both the EI and the RI, suggesting a possible link between perceptual and motor timing precision. We tested the association between α-β PAC in EI and RI and behavioral precision measured by invCV (duration production / standard deviation, see Methods). We choose invCV as opposed to CV for its convenient positive association with precision, which facilitates interpretation of regression results in relation to α-β PAC. invCV in a straightforward manner captures the width of a distribution of time reproduction. A regression model using α-β PAC to predict invCV showed no significant effects in EI [all p > 0.05] or in the RI [p > 0.1].

Alternatively, the precision of timing in this behavioral paradigm may depend on the relative α-β PAC in the EI and RI: for example, if the temporal goal was noisily encoded during the EI, its maintenance in RI may not yield precise temporal reproduction. Therefore, a larger PAC in EI may support higher precision. To test this, we used the difference between the strength in α-β PAC in EI and RI (EI MI_α-β_ (EI) – MI_α-β_ (RI) as a predictor for invCV. The regression model, using the difference in α-β MI between the initial bin in EI and in RI, showed that the strength of α-β oscillatory coupling significantly predicted the behavioral invCV [F(172) = 2.8, P = 0.006; Fig. 5ab]. To ensure that the effect was stable over time, we then tested the difference of α-β coupling strength between the last bin in EI and the first bin in RI: the regression model showed that the strength of α-β oscillatory coupling significantly predicted the behavioral invCV [F(179) = 2.5, P = 0.014; Supp Fig. 1ab]. The analysis, based on Akaike Information Criterion (AIC, Wagenmakers, 2014), showed that both models containing the difference of α-β PAC strength between RI and EI as predictors were justified as compared to the model including only the intercept [ΔAIC = −5.5, P = 0.006; ΔAIC = 4.0, P = 0.014; respectively].

**Figure 5.**
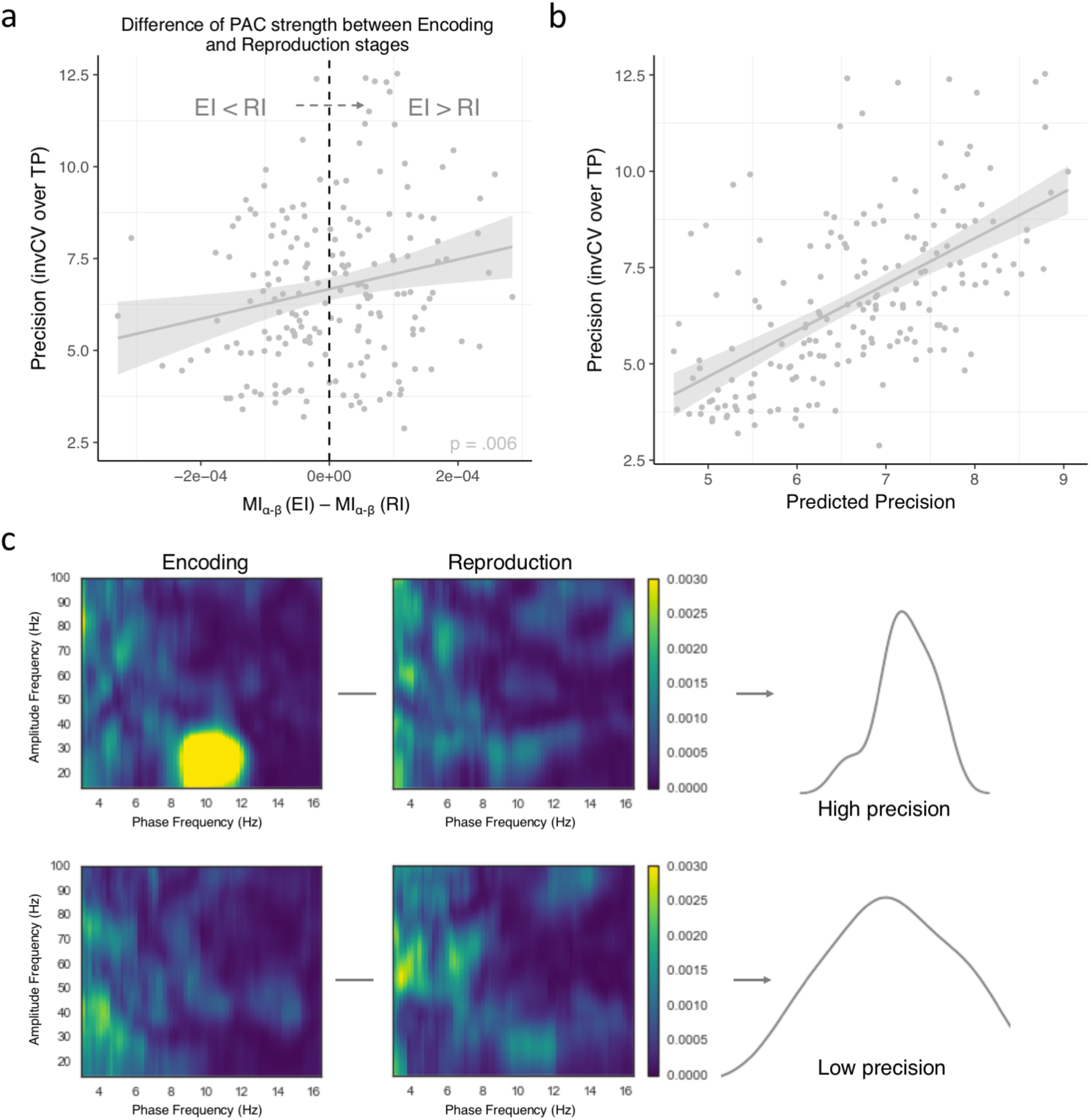
The difference α-β coupling strength between encoding and reproduction informs on the precision of internal timing. a. α-β PAC difference between Bin 1 of the EI and Bin 1 of the RI was significantly correlated with the precision of temporal reproduction (invCV). The model outcomes are also plotted in panel b. b. The fitted values of precision are plotted against the precision values predicted by the mixed model, informed by the difference of α-β PAC strength. c. Two examples ofthe effect plotted in panels a and b. Comodulograms in EI and RI (left and right panels,) for two subsets of trials (rows) with high reproduction precision (top) and with low reproduction precision (bottom). The subset with a higher behavioral precision is shown on the top row and displays larger α-β PAC in EI than in RI.

In both models containing the difference between EI and RI, the α-β PAC strength as predictor in addition to their interaction with duration was not warranted [ΔAIC = −1.7, P > 0.1; ΔAIC = −1.6, P > 0.1; the first and last bin respectively], indicating that the association between precision and EI-RI α-β PAC strength did not depend on the duration being reproduced.

Previous studies investigating phase-amplitude coupling have indicated that MI estimation could be confounded by the estimation of phase and power frequencies (e.g. Aru et al., 2016). To assess whether the association between PAC strength and behavioral invCV was exclusively driven by oscillatory coupling, and not confounded by the power in α or β bands, we tested whether the inclusion of α or β power was justified in the model predicting invCV. Similarly, to α-β PAC, the α power, β power were computed as a difference between EI and RI. The inclusion of α, β power, were not justified in the model predicting invCV for the first bin [ΔAIC = −0.7, P > 0.1; ΔAIC = −1.8, P > 0.1; respectively], and the last bin [ΔAIC = −1.1, P > 0.1; ΔAIC = −1.8, P > 0.1; respectively].

## Discussion

We investigated the role of phase-amplitude coupling in the encoding and in the reproduction of time intervals. Our present results provide evidence that the strength of α-β PAC leverages the precision of timing, extending previous observations in a temporal production task (Grabot, Kononowicz et al., 2017). Here, we specifically show the stable implication of α-β PAC during the encoding of a duration and during the reproduction of the encoded temporal interval. We also show the stability of this effect across three different supra-second durations. Crucially, we found that the difference in α-β PAC between the encoding and the reproduction of temporal intervals was associated with the precision of reproduced intervals (Fig. 5c). This result showed that behavioral precision was not solely a function of α-β PAC strength in the reproduction stage, but rather incorporated a code for the representation of duration encoded earlier in perception, was maintained until motor execution. With this study, we thus provide additional evidence that the strength of α-β PAC indexes the precision with which temporal information is maintained in the brain.

### α-β PAC maintains the precision of temporal representations

Previous work suggested an important role of β oscillations in temporal cognition: β oscillations have been observed during time production and temporal expectations, suggesting that β oscillations may index a neural code for time estimation. During motor timing (Kononowicz & Van Rijn, 2015; Kononowicz et al., 2017), an increase in β power indexes the length of produced time intervals. This pattern has also been reported during perceptual timing (Kulashekhar et al., 2016) and shown to be causally related to time estimation (Wiener et al., 2017). β oscillations have also been suggested to control temporal predictions (Arnal et al., 2014; Fujioka et al., 2012; Teki & Kononowicz, 2016; Iversen et al., 2009). Altogether, prior results implicate β activity in coding duration.

On the other hand, α oscillations and PAC have been implicated in the maintenance of task-relevant information in working memory (Roux & Uhlhaas, 2014). In line with this view and with recent work (Grabot, Kononowicz et al., 2017), α-β PAC was interpreted as reflecting the active maintenance (α) in the timing network of a neural code for duration (β), thereby controlling the precision of timing information. This precision hypothesis predicted an association between α-β PAC and behavioral precision to occur not solely in the RI but also in any period of time during which maintenance of temporal precision would be needed. The current results support this hypothesis considering that behavioral precision was best predicted by the α-β PAC difference between the RI and EI. More specifically, behavioral precision was a function of the strength of PAC during the encoding stage relative to the strength of PAC in the reproduction stage.

This indicate that if the temporal goal is encoded with a limited degree of precision in the EI, its maintenance in RI would not lead to precise temporal reproduction: to the contrary, stronger PAC in the EI did support higher precision of reproduced durations. This imbalance suggests that α-β PAC primarily supports the encoding of a temporal target as opposed to the sole precision of motor commands during temporal production. In line with the precision hypothesis, α-β PAC appears to index the maintenance of higher level temporal representations distinct from the control of motor commands.

Here, the data do not support evidence for the motor control hypothesis. The motor control hypothesis relies on the observation that β oscillations are strongly associated with motor functions, such as movement preparation (Kilavik et al., 2013) and motor control (Engel & Fries, 2010). For example, alternating beta oscillations lead to changes in movement speed (Pogosyan et al., 2009) and successful withholding from movement (Swann et al., 2009). Additionally, Tzagarakis et al., (2010) showed that β amplitude is modulated by the amount of information constraining future movement parameters. Together, these results suggest that beta oscillations carry a multitude of movement parameters. In this context α-β PAC could be seen as neural code that maintains the multitude of movement parameters, where β oscillations code for movement parameters and precision of motor execution is one of the dimensions in the parameter space. The motor control hypothesis predicts that α-β PAC would be only observed in the RI. Secondly, the association between α-β PAC and behavioral precision would only be observed in the RI. However, neither of the predictions of the motor control hypothesis was the case in the current study as we found clear α-β PAC in the EI and the difference in α-β PAC between encoding and reproduction of temporal intervals was associated with the precision of reproduced intervals.

Taken together, these results strengthen the suggestion that α-β PAC in timing and motor timing is not just a function of motor control precision or precision of motor commands. Instead, the obtained results are in line with the working memory hypothesis that offer the view of α-β coupling supporting endogenous maintenance of temporal goal and as such could represent a neural code supporting working memory for duration.

### α-β PAC may support working memory for time

Working memory is a capacity for storing and manipulating information (Baddeley, 1992) and has been broadly studied in many domains (Baddeley, 2012). It is far from clear how working memory extends to interval timing (Matell et al. 2005). Particularly, time reproduction tasks may strongly rely on working memory capacity as suggested by correlational psychophysical studies (Baudouin et al., 2005). This suggests that working memory in the current study may be highly implicated in time keeping function.

From the neuroimaging perspective, a large body of work indicated that maintenance of information in working-memory is supported by PAC in the θ-γ regime (Axmacher et al., 2010; Fell & Axmacher, 2011; Alekseichuk et al., 2016; Leszczynski et al., 2015). For example, enhanced PAC has been associated with increased working memory load (Axmacher et al., 2012) and patients with stronger PAC were capable of maintaining longer sequences in working memory (Leszczynski et al., 2015), suggesting a neural syntax for underlying information maintenance in working memory (Lisman & Jensen, 2013). Along these lines, a recent model proposes a link between working memory supported by θ-γ regime and the retrieval of information during interval timing (Gu et al., 2015; 2018), such that interval timing and working memory differ in terms of which oscillatory features of the oscillating network is used for the extraction of relevant temporal features. At the same time, the oscillatory framework by Gu et al., (2015) has a certain degree of flexibility with respect to the range of frequencies it can accommodate (Teki et al., 2016). Nonetheless, assuming that working memory in interval timing would involve collection of a discrete units of information (van Wassenhove, 2016) as in classical working memory tasks (Siegel et al., 2009), one could expect to see θ-γ coupling at least in the encoding stage, during which participants encoded the duration between two short tones. Despite the extended period of temporal information encoding that was not confounded by other factors such as motor preparation, we found no compelling evidence for the utilization of θ-γ coupling in duration encoding. Although we did not find a typical θ-γ coupling, we did observe α-β coupling, which characteristics did not conform to predicted memory mechanisms. It is also important to note that some studies have previously reported interesting characteristics for β oscillations, which followed parametric modulations during cued working memory when quantitative magnitudes had to be maintained (Spitzer & Blankenburg, 2011). Importantly, duration was one of the tested magnitudes (Spitzer et al., 2014), suggesting that α-β coupling could support some aspects of working memory for duration. Future studies could investigate whether α-β coupling extends the precision with which we encode other magnitudes and what role α-β coupling play in the delay period between encoding and reproduction interval.

### α-β PAC in explicit and implicit timing

To further understand the specificity of representations coded by α-β coupling, future studies could investigate α-β coupling in different timing contexts. For example, in explicit and implicit timing, in which the key distinction would be whether task instructions require participants to provide an explicit estimate of duration or not (Coull & Nobre, 2008; Herbst & Obleser, 2017). In current and previous work (Grabot, Kononowicz et al., 2017), α-β PAC was found during explicit timing tasks. Yet, if such coupling regime extends to implicit timing (e.g., foreperiod paradigm; Praamstra, 2010), temporal expectation (Breska & Deouell, 2014; Nobre et al., 2007), the precision hypothesis could reflect a generic property of the PAC strength indexing the temporal precision of information maintenance in brain networks. There is emerging evidence that delta-beta PAC (θ-β PAC) is sensitive to the occurrence, or absence, of a predicted external event in time (Cravo et al., 2011). θ-β PAC has been found to increase in response to predictable cues in temporal orienting (Mento et al., 2018). Together, these results suggest that at least some form of PAC supports anticipatory processes in implicit timing. Of course, anticipation like processes can play a role in explicit timing. Therefore, the future studies should explore functionalities of coupling regimes jointly in explicit and implicit timing.

## Conclusion

We report that α-β PAC is present during the encoding and during the reproduction of time intervals. Timing precision results from the balance between the strength of α-β PAC during the encoding and during the reproduction of intervals. We suggest that α-β PAC reflects the precision of temporal representations in the human brain.

---

**Supplementary Figure 1.**
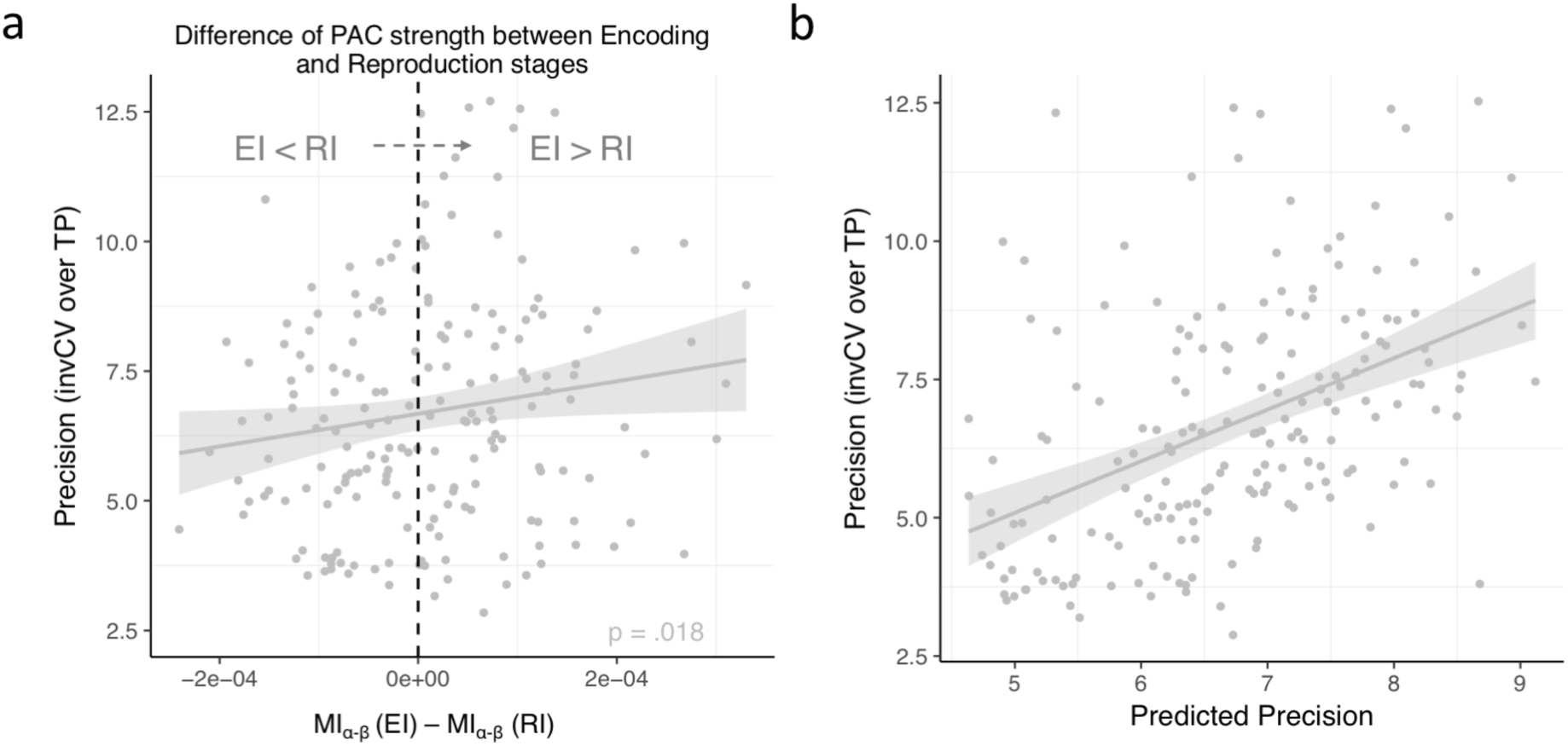
α-β PAC difference between reproduction and encoding indexes the precision of temporal reproduction. a. α-β PAC difference between the last bin of the EI and Bin 1 of the RI were significantly correlated with precision of temporal reproduction (invCV). Model outcomes are also plotted in panel ‘b’. b. The fitted values of precision are plotted against the precision values predicted by the mixed model relying on the difference in the PAC strength.

